# The linkage of methionine addiction, overmethylation of histone H3 lysines and malignancy demonstrated when cancer cells revert to methionine-independence

**DOI:** 10.1101/2020.12.04.412437

**Authors:** Jun Yamamoto, Sachiko Inubushi, Qinghong Han, Yoshihiko Tashiro, Norihiko Sugisawa, Kazuyuki Hamada, Yusuke Aoki, Kentaro Miyake, Ryusei Matsuyama, Michael Bouvet, Steven G. Clarke, Itaru Endo, Robert M. Hoffman

## Abstract

Methionine addiction is a fundamental and general hallmark of cancer and is an area of current intense interest. Methionine addiction results from the overuse of methionine by cancer cells for excess transmethylation reactions. In order to identify excess transmethylation reactions in cancer and further understand the basis of methionine addiction, we compared the histone H3 lysine-methylation status and malignancy between methionine-addicted cancer cells and their methionine-independent revertants which have regained the ability to grow on low levels of methionine or independently of exogenous methionine. The levels of trimethylated histone H3 lysine marks were reduced in methionine-independent revertants compared to parental cancer cells *in vitro*. Tumorigenicity and experimental metastatic potential in nude mice were also highly reduced in the methionine-independent revertants compared to the parental cells. Our present results demonstrate that overmethylation of histone H3 lysines is linked with methionine addiction of cancer and to malignancy which suggests a possible causal relationship.

## Introduction

Methionine addiction is a fundamental and general hallmark of cancer discovered by one of us (RMH) ^1–4^ and is an area of current intense interest ^5–7^. Methionine addiction is characterized by a requirement for exogenous methionine for growth by cancer cells even though the methionine-addicted cancer cells synthesize normal or excess amounts of methionine ^1–4^. Methionine restriction (MR) by either methionine-free medium ^8^ or by a low-methionine diet ^9^ or by methioninase ^10^ selectively arrests cancer cells in the late S/G_2_ phase of the cell cycle, but not normal cells, where the cancer cells become sensitive to cytotoxic chemotherapy^11^. Methionine addiction results from the overuse of methionine by cancer cells for excess transmethylation reactions^12^, termed the Hoffman-effect, analogous to the Warburg effect for glucose overuse by cancer cells ^13,14^. Methionine addiction is tightly linked to other hallmarks of cancer ^13,15^ and is thought possibly the very basis of malignancy itself ^16^.

Although we have long since known methionine is overused for transmethylation reactions cancer cells ^12,17^, we have poorly understood the fate of at least a significant amount of the excess methyl groups that were transferred. Our previous studies have shown that histone H3 lysine marks are overmethylated in cancer cells compared to normal cells^18,19^ and that the histone H3 lysine overmethylation is unstable during MR of methionine-addicted cancer cells which arrests their proliferation. In contrast the lesser amount of histone H3 lysine methylation is stable in normal cells under MR, which unlike cancer cells do not arrest their proliferation^18^. These results suggested that histone H3 lysine overmethylation may be related to methionine addiction.

In the present study, we report that methionine-independent revertants, selected in methionine-free medium containing homocysteine or low methionine from methionine-addicted cancer cells, lose over-methylation at histone H3 lysine and malignancy compared to methionine-addicted cancer cells. Importantly, methionine-independent revertants lose their requirement for exogenous methionine. Our results indicate that histone H3 lysine overmethylation is linked with methionine addiction and malignancy, suggesting a possible causal relationship.

## Results

### Cancer cells selected to grow in low-methionine media revert from methionine-addiction to methionine independence

We used recombinant methioninase (rMETase) to deplete methionine *in vitro*. The level of methionine in the medium was rapidly decreased to less than 30% within 1 hour after addition of 1 U/ml of rMETase (Fig. 1B).

**Figure 1.**
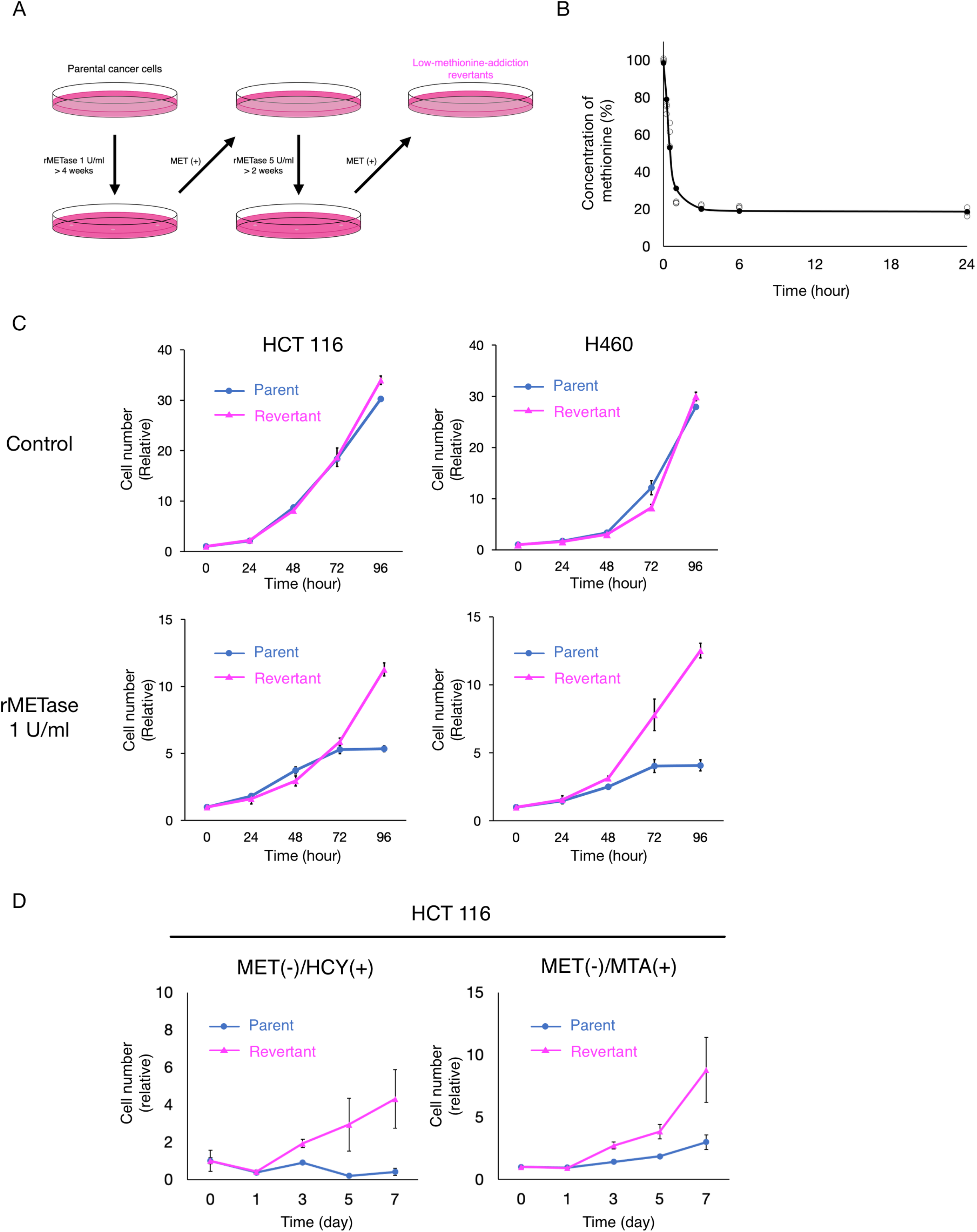
Comparison of sensitivity to methioninase of methionine-addicted cancer cells and methionine-independent revertants *in vitro*. (A) Diagram of the establishment of low-methionine-requirement revertants. (B) Recombinant methioninase (rMETase) depletes methionine in cell culture medium. rMETase was added to the medium (1 U/ml) and the methionine levels in the medium were measured at 0 hour, 15 minutes, 30 minutes, 1 hour, 3 hours, 6 hours and 24 hours (n = 3). (C) Cell proliferation assay of methionine-addicted parental cancer cells and their methionine-independent revertants under methionine restriction. Cells were cultured in methionine-containing medium or methionine-containing medium with rMETase (1 U/ml) for 24, 48, 72 and 96 hours (mean ± SEM, n = 3). (D) Cell proliferation assay of methionine-addicted parental cancer cells and their methionine-independent revertants in methionine-free medium containing 200 μ M DL-homocysteine and 250 nM methylthioadenosine (MTA) with 10% dialyzed fetal bovine serum.

To compare the methionine requirement of methionine-addicted parental cancer cells and low-methionine-requirement revertants derived from the parental cells, we then evaluated their cell proliferation kinetics under methionine restriction effected by rMETase. The proliferation of parental methionine-addicted cancer cells arrested within 72-96 hours of methionine restriction. In contrast, methionine-low-requirement revertants were able to continuously proliferate under methionine restriction, similar to normal cells^13,15,17,20^ (Fig. 1C). We also evaluated the cell proliferation of low methionine-requirement revertatnts in methionine-free medium containing homocysteine or methylthioadenosine (MTA). Low-methionine-requirement revertants proliferated in medium which methionine was replaced by either homocysteine or MTA similar to normal cells (Fig. 1D). In contrast, parental methionine-addicted cancer cells arrested in either replacement medium. These results indicate that the revertants have lost their methionine addiction and requirement for exogenous methionine.

### Methionine synthase and methylthioadenosine phosphorylase are upregulated in methionine-independent revertants

To determine the differences in methionine metabolism in methionine-addicted cancer cells and methionine-independent revertants, we examined changes in protein expression levels for enzymes involved in methionine biosynthesis in parental methionine-addicted HCT 116 and H460 cells and their respective methionine-independent revertants (Fig. 2A). We found the methionine synthase (MTR) was modestly upregulated in methionine-independent revertants of each cell line and that methylthioadenosine phosphorylase (MTAP) was strongly upregulated in the HCT 116 revertant cells and moderately upregulated in the H460 revertant cells (Fig. 2B).

**Figure 2.**
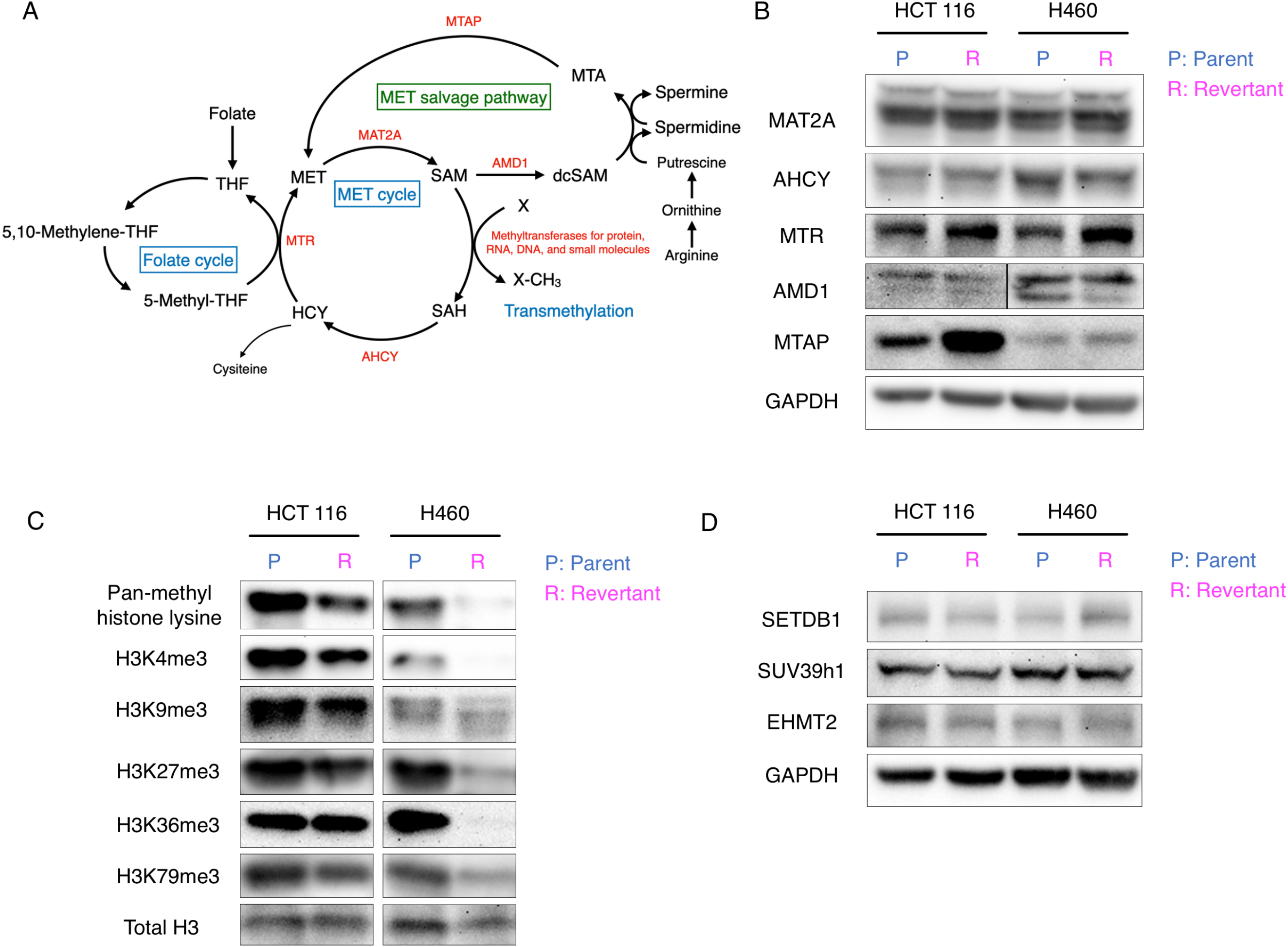
Methionine and methyl group metabolism in parental methionine-addicted cancer cells and their methionine-independent revertants *in vitro*. (A) The pathways of methionine metabolism. (B) Immunoblot of the enzymes related to methionine metabolism. (C) Immunoblot of pan-methyl H3 lysine methylation and trimethylated histone H3 lysine marks in methionine-addicted cancer cells and methionine-independent revertants grown in methionine-containing medium. (D) Immunoblot of H3K9 methyltransferases in methionine-addicted cancer cells and methionine-independent revertants grown in methionine-containing medium.

### Overmethylation of histone H3 lysine marks is strongly decreased in methionine-independent revertants

We then compared the methylation status of histone H3 lysine marks in methionine-addicted parental cells and methionine-independent revertants. Parental cancer cells and methionine-independent revertants were cultured in normal methionine-containing medium for 96 hours and histones were extracted. The overall level of lysine methylation of histone H3 was decreased in the methionine-independent revertants compared to parental methionine-addicted cells even in the presence of methionine (Fig. 2C). The levels of specific methylation marks including H3K4me3, H3K9me3, H3K27me3, H3K36me3 and H3K79me3 were also decreased in the methionine-independent revertants compared to parental methionine-addicted cells in the presence of the methionine (Fig. 2C). The levels of pan-methyl lysine of H3, H3K4me3, H3K9me3, H3K27me3, H3K36me3 and H3K79me3 were much lower in HCT 116-R cells than in HCT 116 cells as shown by immunoblotting(Fig. 2C). However, the expression levels of several histone methyltransferase enzymes which methylates H3K9me3 did not change in the methionine-independent revertants (Fig. 2D).

These results show that methionine addiction and histone H3 lysine-mark overmethylation are linked to decreased use of methionine for histone H3 lysine methylation.

### Methionine-independent revertants lose malignancy

To compare the malignancy of parental methionine-addicted cancer cells and methionine-independent revertants derived from them, the tumorigenicity of the parental and revertant cells was compared in subcutaneous xenograft mouse models. Methionine-addicted parental HCT 116 cells and their methionine-independent revertants (HCT 116-R) were compared for their ability to form tumors in nude mice. The mean tumor volume was significantly lower in HCT 116-R tumors than HCT 116 tumors after injection of 1 × 10^6^ cells in nude mice (P = 0.0085), and only half of 10 mice formed tumors in HCT 116-R compared to all mice with HCT 116 (Fig. 3A). While 8 out of 10 mice injected with 5 × 10^5^ parental methionine-addicted HCT 116 cells formed tumors, no mice injected with 5 × 10^5^ HCT 116-R cells formed tumors.

**Figure 3.**
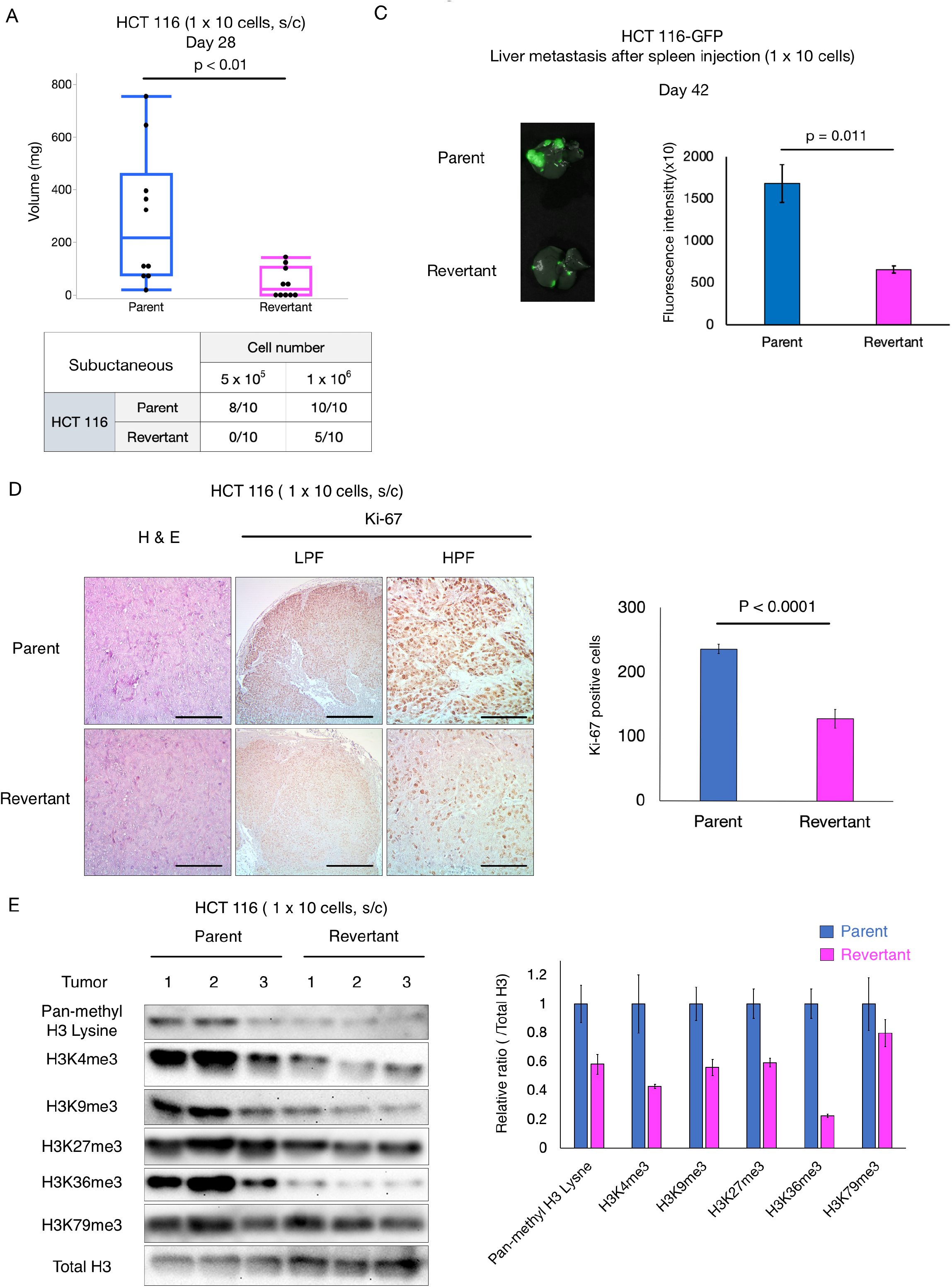
Comparison of malignancy of methionine-addicted cancer cells and their methionine independent revertants *in vivo*. (A) Mean tumor volume of HCT 116 and HCT 116-R at 28 days after 1 × 10^6^ cells were injected (n = 10. *, p < 0.01). (B) Number of the tumors in nude mice at day 28 of HCT 116 and HCT 116-R. (C) Experimental liver metastasis: parental methionine-addicted HCT 116-GFP and methionine-independent revertants HCT 116-R-GFP. Right representative fluorescence image of liver metastasis. Left Fluorescence density (mean ± SEM, n = 3). (D) Left: Representative images of H & E staining and immunohistochemical staining for Ki-67 of HCT 116 and HCT 116-R subcutaneous tumors. LPF: low power field (40×). HPF: high power field (200×). Scale bar: 100 μm (H & E and HPF of Ki-67), 500 μm (LPF of Ki-67). Right: Quantification of the Ki-67 positive cells. Five high-power fields were randomly chosen (400×) and positive cells compared (mean ± SEM, n = 4). *, p < 0.0001. (E) Left: Immunoblot of pan-methyl H3 lysine and trimethyl histone H3 marks in subcutaneous tumors formed from HCT 116 and HCT 116-R (n = 3). Right: The ratio of pan-methyl H3 lysine or trimethyl-histone H3 marks / total H3 in the tumors formed from HCT 116 and HCT 116-R (mean ± SEM, n = 3).

The metastatic potential of HCT 116-GFP expressing cells and HCT116-R-GFP expressing cells was also compared after spleen injection the two cell types in nude mice. The parental HCT 116-GFP cells formed significantly more experimental liver metastasis compared to revertant HCT-116-R-GFP cells (p = 0.011) (Fig. 3B). These results show that methionine-independent revertants have significantly reduced malignancy.

Immunohistochemistry staining showed that the number of Ki-67-positive cells was significantly lower in the HCT 116-R subcutaneous tumors compared to the parental HCT 116 subcutaneous tumors (n = 4, P < 0.0001) indicating reduced in vivo proliferation of the methionine-independent revertants (Fig. 3C). The tumorigenicity, metastasis and cell proliferation results indicate that methionine-independent revertants are much less malignant than their methionine-addicted parents.

The malignancy decreases in methionine-independent revertants compared to parental methionine-addicted cancer cells is linked with a decrease in histone H3 lysine methylation as described above. The results of the present study and our previous study ^21^ suggest that overmethylation of histone H3 lysine marks linked to both methionine-addiction and malignancy and their relationship may be causal.

### RNA sequencing of methionine-addicted parent and methionine-independent revertant colon cancer and lung cancer cell lines show no large scale difference in gene expression

To determine if large-scale changes in gene expression are involved in the reversion of cancer cell lines from methionine addiction to methionine-independence and loss of tumorigenicity, we compared mRNA transcript levels in parent and revertant cells from the H4CT 116 cell line and the H460 cell line. In Table 1, we show genes with transcription levels that were significantly elevated or depressed more than two-fold between the parent and revertant cell lines. Of the some 19,000 human genes measured for expression, only a small number met these criteria. For the HCT 116 parental cells, we found 14 down-regulated at least two-fold and 7 genes that were up-regulated by at least two-fold compared to HCT 116 revertant cells (Table 1A). The corresponding numbers for the H460 cells were 80 genes down-regulated and 14 up-regulated genes in the revertants compared to parental cells (Table 1B). We found no common genes whose expression changed two fold or more between the lung-cancer and colon-cancer derived cell lines. These results suggest that there may be no common transcriptional pathway that would account for the reversion of methionine-addicted cancer cells to methionine-independence, or the methods used were not sensitive enough to detect such common genes. With the possible exception of the FOLR1 gene encoding the alpha folate receptor, none of these genes with a two-fold or more change appeared to be involved in methionine metabolism.

**Table 1.**
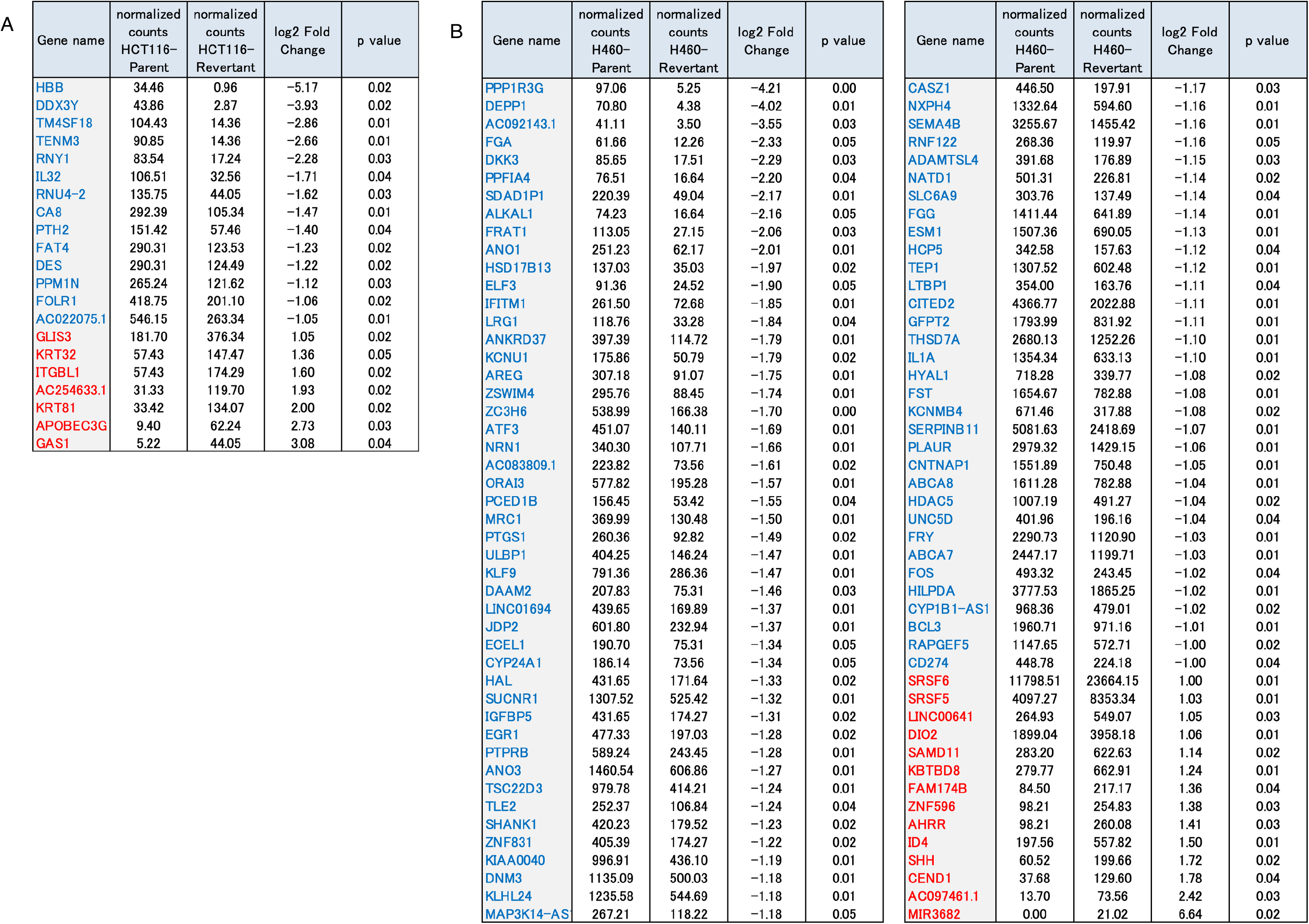
Genes with expression levels that are significantly different between methionine-addicted cancer cells and methionine-independent revertants.

### Changes in expression of methionine metabolism genes in methionine-addicted parental cancer cells and methionine-independent revertants

We then examined the changes in gene expression for genes involved in the biosynthesis of methionine, its conversion to *S*-adenosylmethionine, and the use of *S*-adenosylmethionine for polyamine biosynthesis and transmethylation reactions, as well as genes encoding enzymes involved in demethylation (Figure 2B; Table 2; Supplemental Table 1). These genes included methionine synthase and related proteins, *S*-adenosylmethionine synthetases, enzymes and proteins of the folate cycle that provide methyl groups for methionine biosynthesis, methionine salvage and polyamine biosynthetic proteins, and enzymes involved in the metabolism of the product of all methyltransferases, *S*-adenosylhomocysteine. Additionally, we examined the expression of genes for 182 putative and established *S*-adenosylmethionine-dependent methyltransferases, catalyzing methylation of protein, RNA, DNA, and small molecules as well as genes for 17 putative and established demethylases of protein, DNA, and RNA. We found very few changes in the expression of these genes when we compared the parents and revertants of the HCT 116 and H460 cancer cells. The most interesting statistically-significant change was an almost two-fold increase in the expression of the MAT2A gene encoding the catalytic subunit of the S-adenosylmethionine synthetase in the H460 revertant (p = 0.01) (Table 2A). However, the expression of this gene was decreased in the HCT 116 revertant cells. These results suggest that there are multiple pathways that can lead to reversion to methionine-independence. Additionally, we should point out that changes in mRNA levels need not necessary be correlated with changes in the activities of the protein, since translational and posttranslational controls may play larger roles than transcriptional controls in determining the activities of some enzymes. For example, when we looked at the protein levels of MAT2A by immunoblotting in Figure 2B, we observed no change in the methionine-independent revertants of either of the methionine-addicted cancer cell lines.

**Table 2.**
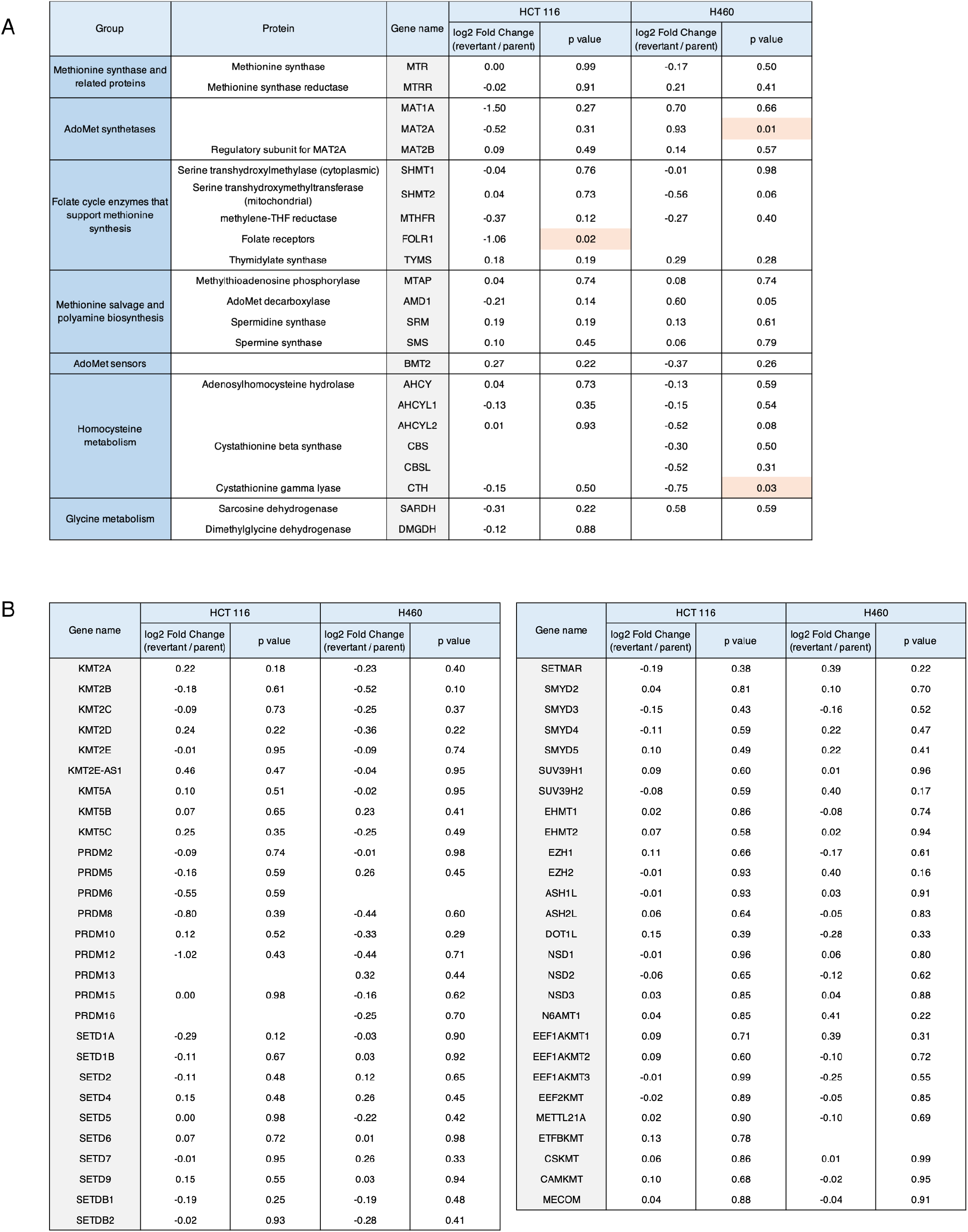
Expression of genes related to methionine metabolism and protein methyltransferases in methionine-addicted cancer cells and their methionine-independent revertants.

As noted above, we also found a statistically-significant change in the expression of the gene for the alpha folate receptor in the HCT 116 revertants (~two-fold decrease; p = 0.02) and the gene encoding an enzyme in the trans-sulfuration pathway from homocysteine to cysteine (cystathionine gamma lyase, which had a 1.7 fold decrease in the H460 revertants; p = 0.03). However, neither change was common to the revertants of both colon-cancer and lung-cancer cell lines (Supplemental Table 1).

We noted statistically-significant decreases in the expression of the genes in the revertant lines from both the HCT 116 and H460 cells for two protein lysine demethylases, KDM6B in the H460 revertants (1.9 fold, p = 0.04) and KDM7A in the HCT 116 revertants (1.5 fold; p = 0.02) (Table 2B). However, from the immunoblotting results in Figure 2D, we observed decreases in the level of histone H3 trimethylation at this site in both the colon cancer and lung cancer revertnants, indicating that the decrease in levels of histone H3 lysine methylation in the methionine-independent revertant is not due to demethylases but possibly to decreased *S*-adenosylmethioinine utilization.

The present results suggest that there are few significant changes in gene expression and associated with reversion to a methionine-independent phenotype

## Discussion

Methionine dependence of cancer has been known since 1959 when Sugimura showed that tumors in rats slowed their growth when methionine was removed from their diet compared to when other amino acids were removed from the diet ^22^. It was not until the early 1970s that methionine dependence was found in cultured cancer cells ^23,24^ and it was initially claimed that methionine dependence was due to reduced ability of the cancer cells to synthesize methionine from homocysteine 24. At that time in 1976, one of us (RMH) demonstrated that cultured methionine-dependent cancer cells made normal or more-than-normal amounts of methionine from homocysteine, but still required exogenous methionine in order to proliferate ^1^, first demonstrating that cancer cells were methionine addicted. The requirement for exogenous methionine by methionine-addicted cancer cells was later confirmed by Tisdale in 1984 25 and in 2019 by Tam’s group ^7^. It is important to note that we observed that as little as 1μM methionine stimulated the growth of methionine-addicted cancer cells to initiate rapid proliferation on methionine-free homocysteine-containing medium in which they were arrested. ^8^. These results give rise to speculation that exogenous methionine may have a special use in methionine-addicted cancer cells. Future metabolomics experiments may give us more information on the fate of exogenous methionine in methionine-addicted cancer cells compared to methionine-independent revertants.

We previously showed that an increase in the rate of synthesis of methionine from homocysteine was not necessary for reversion to methionine independence^20^, further demonstrating that decreased methionine synthetic capacity was not the basis of methionine addiction. Then we found that despite the large amounts of endogenous methionine made by cancer cells, under restriction of exogenous methionine the cancer cells had very low levels of free methionine, low levels of *S*-adenosylmethionine (SAM) and a low ratio of SAM to *S*-adenosylhomocysteine (SAH) compared to normal cells and methionine-independent revertants^1–4,15,20^. This was explained next when we observed that all cancer cells tested had elevated rates of transmethylation compared to normal cells which used excess amounts of methionine and SAM and depleted their cellular pools ^2,3,12^. We then understood that methionine addiction was due to overuse of methionine for transmethylation ^12^. This was confirmed when it was demonstrated that overall transmethylation was strongly reduced in methionine-independent revertants derived from the methionine-addicted cancer cells ^16,17^. The phenomenon of methionine-addicted cancer cells is termed the Hoffman-effect ^14^.

Tam’s group in 2019 ^7^, confirmed our results of 44 years previously ^1^ of high methionine flux in cancer cells, in this case tumor-initiating cancer cells, resulting in a requirement for exogenous methionine for their proliferation. Tam et al. showed abundant levels of trimethyllysine residues in histone H3 in tumor-initiating cells, compared to cancer cells which were not tumor-initiating cells ^7^. However, the levels of trimethyllysine residues in histone H3 in normal cells or methionine-independent revertants were not measured by Tam et al. Tam’s group showed that MR eliminated tumor-initiating cells ^7^, similar to our previous observation that MR eliminated clonogenic cancer cells ^8^. Mentch showed that the level of trimethyllysine residues in histone H3 were unstable in cancer cells under MR. However, the did not examine the stability of these methylated residues in normal cells under MR ^26^. Our recent previous study also demonstrated that histone H3K4me3 and H3K9me3 were unstable in cancer cells under MR; and we showed for the first time H3K4me3 and H3K9me3 were stable in normal cells under MR ^18^, which may explain at least in part why cancer cells arrest under MR and normal cells do not.

And most importantly, the present study shows that all measured trimethylated histone H3 marks (as well as H3 pan methylatlion) show overmethylation in parental methionine-addicted cancer cells but not in methionine-independent revertants that have decreased malignancy and have lost methionine addiction, and can proliferate under MR or in the absence of methionine when it is replaced by homocysteine or MTA. The present study and our recent study ^18^ suggest that malignant cells require histone H3 overmethylation in order to grow *in vitro* and to form tumors and metastasis.

The present study thus shows that pan-overmethylated lysine marks of histone H3 explain at least in part the fate of an important set of methyl-groups in cancer cells and why cancer cells are methionine addicted and not normal cells or methionine-independent revertant cells. Most importantly, overmethylation of histone H3 lysine resdidues is lost along with malignancy and methionine addiction in methionine-independent revertants, derived from methionine-addicted cancer cells. The first methionine-independent revertants isolated 43 years ago from methionine-addicted SV-40-transformed human fibroblasts ^15,20^ and recent methionine-independent revertants of triple negative breast cancer cells had reduced clonogenicity in agar, a surrogate maker of malignancy ^13^, which gave the first hints that methionine addiction may be necessary for malignancy. Our present results therefore further suggest overmethylation of histone H3 lysine marks, including trimethyl histone H3 marks, may be necessary for malignancy as indicated by reduction in tumorigenicity, metastasis and cancer cell proliferation within tumors in methionine-independent revertants along with greatly reduced methylation of histone H3 lysine methylatlion marks.

Although methionine-addicted cancer cells and methionine-independent revertants grew similarly *in vitro* in the presence of methionine, the revertants had greatly reduced malignancy as see by reduced tumorigenicity and metastasis ^27^ and reduced cancer-cell proliferation *in vivo* as shown by the reduced frequency of Ki-67 expression (Fig. 3D). All tested cancer types are methionine addicted as shown *in vitro*, by their much higher sensitivity to methioninase than any tested normal cell type and the cancer cells are selectively arrested by MR ^9,28–34^. The results of the present study indicate that methionine addiction is linked to histone H3 lysine overmethylation and is a general and fundamental hallmark of cancer and is possibly necessary for malignancy.

Methionine restriction alters the expression of certain genes. We and other investigators have previously reported that MR increased TNF-related apoptosis-induced ligand receptor-2 (TRAIL-R2) expression in cancer cells and enhanced the efficacy of TRAIL-R2 targeted therapy ^35,36^. However, the present study shows that there are few significant changes between parental cancer cells and methionine-independent revertants in RNA expression which is surprising given the large reduction of histone H3 lysine marks that are thought to be master regulators of gene expression. These results indicate that the loss of tumorigenicity in methionine-independent revertants may be due to translational and posttranslational controls. Further proteomics studies are needed to investigate translational and posttranslational controls in methionine-addicted cancer cells and their revertants.

In the present study we did not detect a difference of the expression level of histone methyltransferases between methionine-addicted cancer cells and methionine-independent revertants by immunoblotting or RNA expression. In addition, the MAT2A protein level is not changed in the revertants. There are previous studies which report that high expression level of histone lysine methyltransferases and MAT2A are markers of malignancy ^7,37,38^. However, our present study shows that methionine-independent revertants lose malignancy without detected changes in the expression levels of those enzymes.

Also important is the finding by Breillout et al. that as cells become more malignant they become more methionine addicted ^27^. Another important observation is from PET imaging of cancer in patients which consistently shows that [^11^C] methionine gives a much stronger PET signal than [^18^F] deoxyglucose in head-to-head comparisons, demonstrating that cancers are methionine addicted in patients and that the Hoffman-effect is stronger than the Warburg-effect ^39,40^. Other molecules, such as various RNAs, may also be overmethylated in cancer cells ^41,42^. Further experiments are necessary to account for all the transferred methyl groups in cancers and the difference in comparison to normal cells.

In summary, our results suggest that overmethylation of histone H3 lysines is linked with methionine addiction and malignancy of cancer cells, and possibly necessary for both fundamental hallmarks of cancer (Fig. 4). Methionine addiction is being widely recognized as critical for malignancy ^43^ and may provide a universal target for methionine-restriction cancer therapy, such as with methioninase which depletes methionine in tumors rapidly and shown to be broadly effective against all cancer types tested including the clinic ^44^,.

**Fig. 4.**
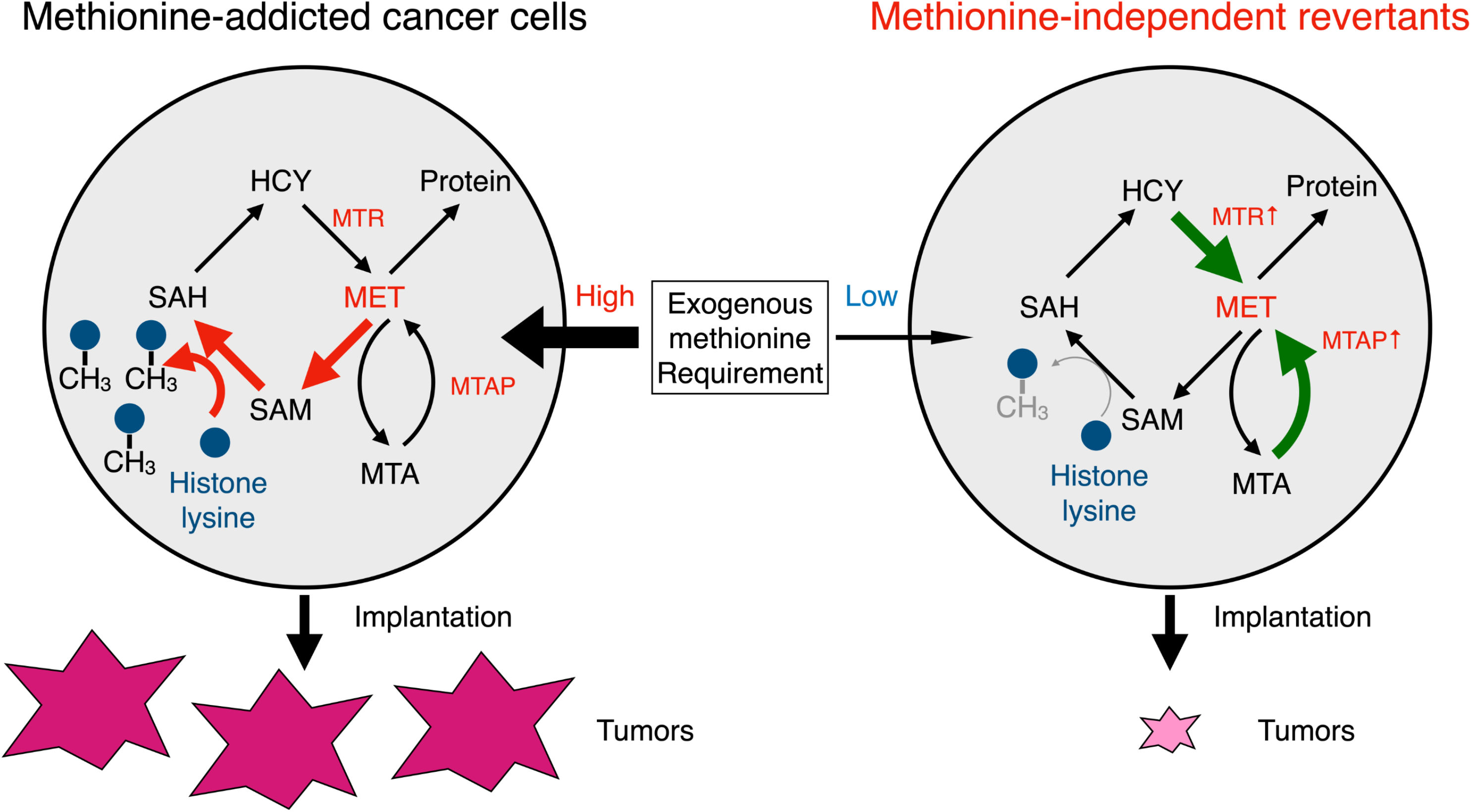

## Materials and Methods

### Cell culture

The HCT 116 human colon cancer cell line and H460 human lung cancer cell line were used in the present study. HCT 116 cells were stably transduced to express green fluorescent protein (GFP) as previously described ^45^. All cell lines were obtained from the American Type Culture Collection (Manassas, VA, USA) and were routinely checked for mycoplasma. Cells were maintained in DMEM supplemented with 10% fetal bovine serum (FBS) and 100 IU/ml penicillin/streptomycin.

### Recombinant methioninase production

Recombinant L-methionine α-deamino-γ-mercapto-methane lyase (rMETase) is a 172-kDa molecular homotetrameric PLP enzyme from *Pseudomonas putida* ^46^. The rMETase gene was cloned in *E.coli*. Fermentation of rMETase-recombinant *E. coli* and purification of rMETase were performed as previously describe ^47^.

### Selection and establishment of methionine-independent revertant cancer cells

The H460 and HCT 116 parental cell lines were cultured in normal methionine-containing medium with 1 U/ml of rMETase for more than one month as a first selection and surviving cells were isolated. These surviving cells were cultured in normal methionine-containing medium with 5 U/ml rMETase for more than 2 weeks as a second selection and surviving cells were isolated. The surviving cells isolated after the second selection were termed H460-R and HCT 116-R respectively.

### rMETase activity assay in vitro

Medium with 1 U/ml of rMETase was incubated, and the methionine levels in the medium were measured with an HPLC (Hitachi L-6200A Intelligent pump; Hitachi, Ltd., Tokyo, Japan) before and after added rMETase as described previously ^48^.

### Efficacy of MR, effected by rMETase, on cell proliferation

Cells were cultured in 96-well plates (1 × 10^3^ cells/well) in normal DMEM overnight. The next day, the medium was changed to normal DMEM or DMEM with rMETase (1 U/ml). Cell proliferation was measured using the Cell Counting Kit-8 (Dojindo, Kumamoto, Japan) after 0, 24, 48, 72 and 96 hours of medium change.

### Cell proliferation of methionine-independent revertatnts compared to parental methionine-addicted cancer cells in methionine-free medium conditioning homocysteine or methylthioadenosine (MTA)

Cells were cultured in 96-well plates (1 × 10^3^ cells/well) in normal DMEM overnight. The next day, the medium was changed to methionine-free medium containing 200 μM DL-homocysteine (Sigma) or 250 nM methylthioadenosine (MTA, Sigma) with 10% dialyzed fetal bovine serum. Cell proliferation was measured using the Cell Counting Kit-8 (Dojindo, Kumamoto, Japan) after 0, 1, 3, 5 and 7 days of medium change.

### Immunoblotting

Extraction of histone and protein from cells and tumors and immunoblotting were performed as described ^18,36^. Antibodies used are listed in Supplemental table 2 Immunoreactive proteins were visualized with Clarity Western ECL Substrate (Bio-Rad Laboratories, Hercules, CA, USA). The UVP ChemStudio (Analytik Jena US LLC, Upland, CA, USA) was used to detect the signals.

### Mouse studies

4-6 weeks old athymic *nu/nu* female mice (AntiCancer Inc, San Diego, CA, USA), were used in this study. All mice were kept in a barrier facility on a high efficacy particulate air (HEPA)-filtered rack under standard conditions of 12 h light/dark cycles. Animal studies were performed with an AntiCancer Institutional Animal Care and Use Committee (IACUC)-protocol specially approved for this study and in accordance with the principles and procedures outlined in the National Institutes of Health Guide for the Care and Use of Animals under Assurance Number A3873-1 ^36^.

### Comparison of *in vivo* tumorgenicity of methionine-addicted parental cancer cells and methionine-independent revertants in a subcutaneous mouse model

Two different doses of HCT 116 and HCT 116-R cells (5 × 10^5^ or 1 × 10^6^ cells / 100 μl PBS each) and were injected subcutaneously into the flanks of nude mice. Each group comprised ten mice. The mice with HCT 116 or HCT 116-R tumors were sacrificed on day 28 and tumor volume was measured at termination^21^.

### Comparison of ability of methionine-addicted parental cancer cells and methionine-independent revertants to form experimental liver-metastasis

HCT 116-GFP and HCT 116-R-GFP (1 × 10^6^ cells / 100 μl PBS) were injected to the spleen of nude mice. Each group comprised three mice. The mice were sacrificed on day 42 and fluorescence intensity was measured at termination with the UVP ChemStudio (Analytik Jena US LLC) ^45^.

### H & E staining and immunohistochemistry

Hematoxylin and eosin (H&E) staining and immunohistochemical staining (IHC) were performed as described^36^. Rabbit polyclonal anti-Ki-67 antibody (1:16,000, 27309-1-AP, Proteintech) was used as a cell proliferation marker. For immunohistological evaluation, two investigators (K.H. and Y.A.) selected the five most abundant microscopic fields of each tissue and counted Ki-67-positive cells (magnification, 400×) in each of the five regions.

### Next generation sequencing

Cells (1 × 10^6^) were pelleted and frozen at −80 °C and provided to GENEWIZ (South Plainfield, NJ, USA) for RNA sequencing. RNA was extracted from the cells and were sequenced by HiSeq sequencing system (Illumina, San Diego, CA, USA).

### Statistical analyses

All statistical analyses were performed with JMP PRO ver. 15.0.0 (SAS Institute, Cary, NC, USA). Student t-test was used to compare between groups for animal studies. Bar graphs show the mean, and error bars express standard error of the mean. A probability value of P < 0.05 is defined as statistically significant.

## Acknowledgements

This work was supported in part by a Yokohama City University research grant “KAMOME Project”. The study was also supported in part by the Robert M. Hoffman Foundation for Cancer Research. Neither organization had a role in the design, execution, interpretation, or writing of the study. This paper is dedicated to the memory of A. R. Moossa, M.D., Sun Lee, M.D., Professor Li Jiaxi and Masaki Kitajima, MD.

## Author Contributions

J.Y., S.G.C and R.M.H designed and performed experiments and wrote the paper; S.I., N.S., K.H., Y.T., Y.A., K.M., R.M., M.B. and S.G.C gave technical support and conceptual advice. Q.H. produced methioninase. Writing, review, and/or revision of the manuscript: J.Y., S.G.C, I.E. and R.M.H.

---

**Supplemental Table 1.**
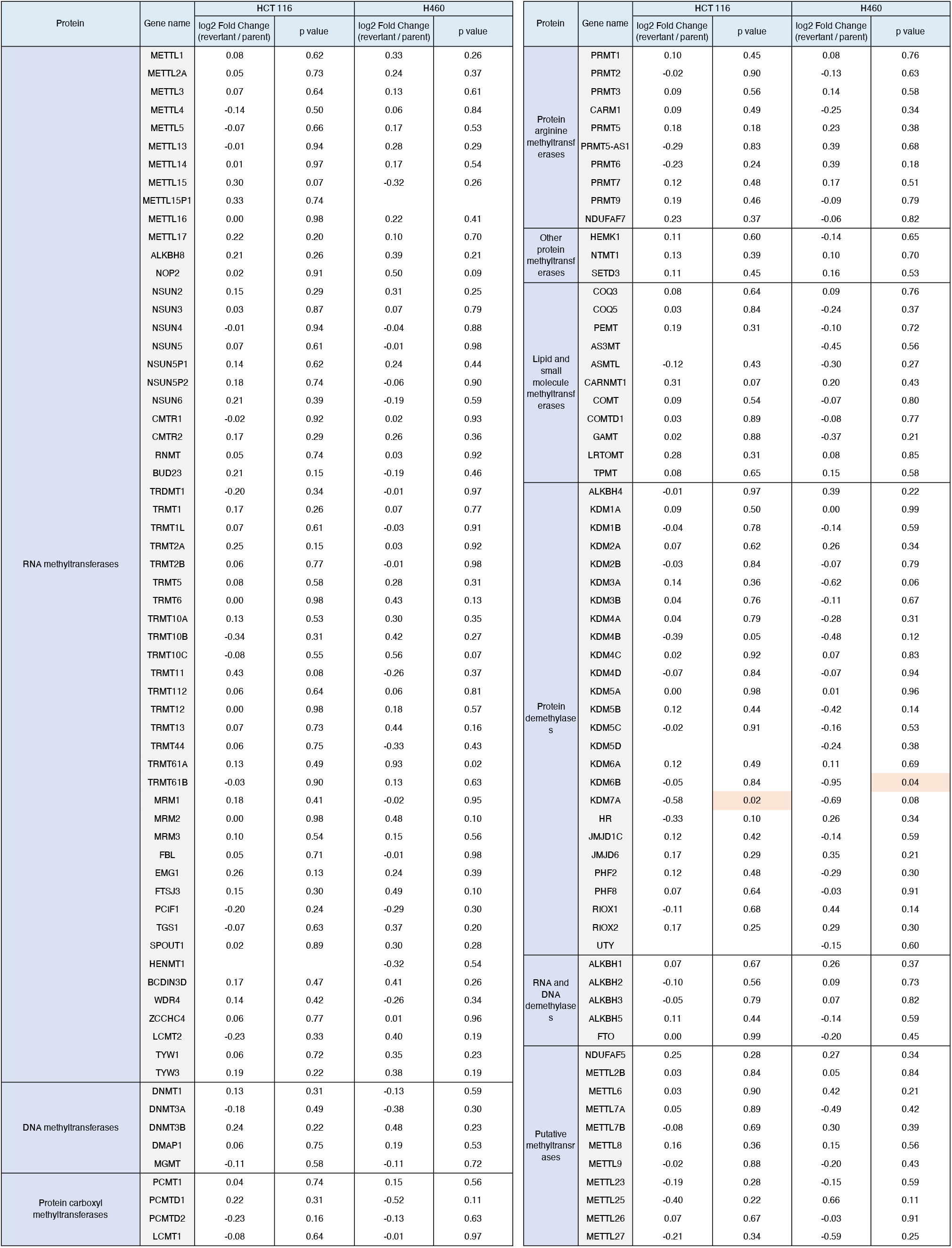
Expression of the genes for methyltransferases, other than protein methyltransferases, in methionine-addicted cancer cells and their methionine-independent revertants.

**Supplemental Table 2.**
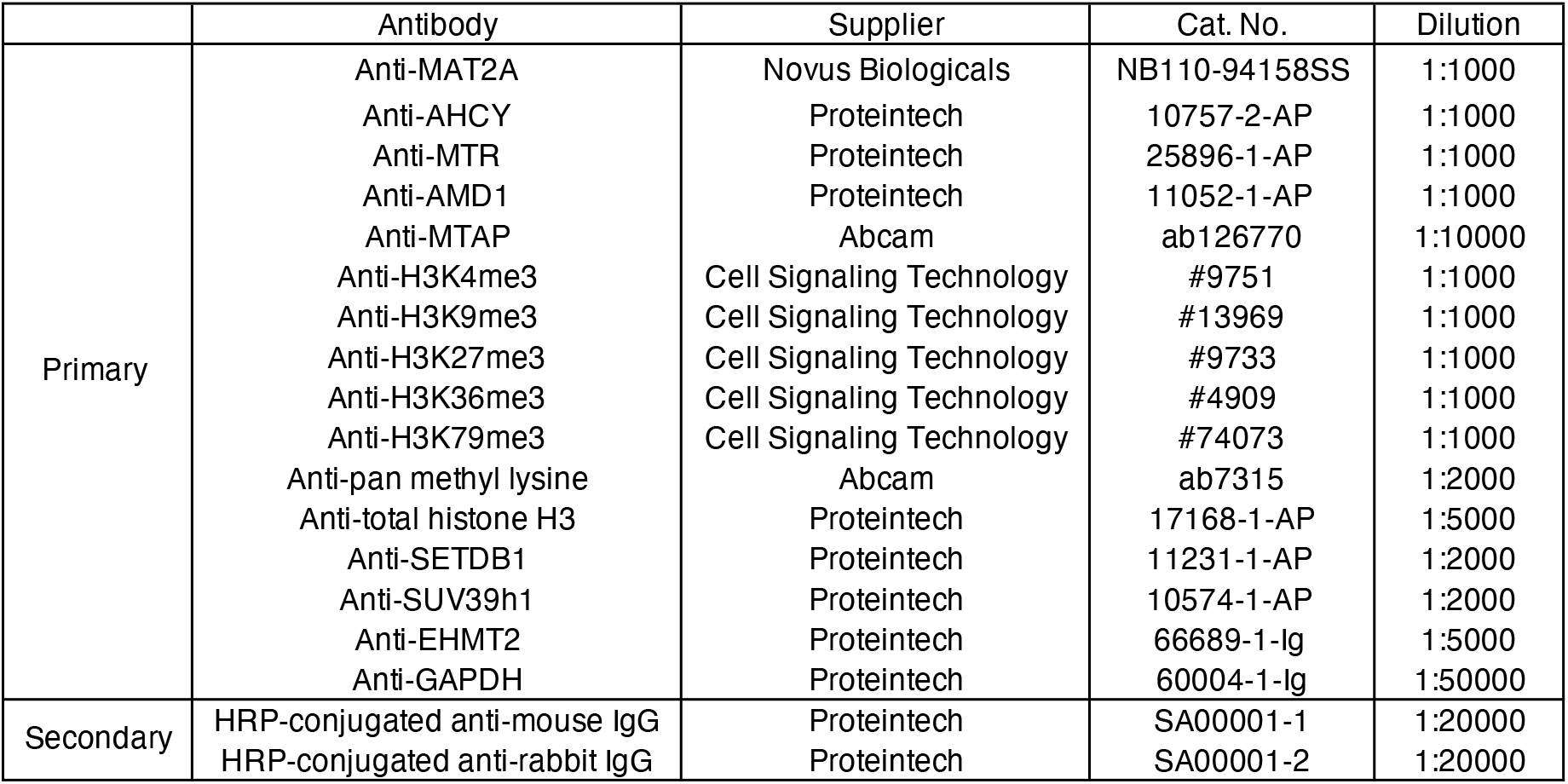
Antibodies used for immunoblotting.

|  | Antibody | Supplier | Cat. No. | Dilution |
| --- | --- | --- | --- | --- |
| Primary | Anti-MAT2A | Novus Biologicals | NB110-94158SS | 1:1000 |
|  | Anti-AHCY | Proteintech | 10757-2-AP | 1:1000 |
|  | Anti-MTR | Proteintech | 25896-1-AP | 1:1000 |
|  | Anti-AMD1 | Proteintech | 11052-1-AP | 1:1000 |
|  | Anti-MTAP | Abcam | ab126770 | 1:10000 |
|  | Anti-H3K4me3 | Cell Signaling Technology | #9751 | 1:1000 |
|  | Anti-H3K9me3 | Cell Signaling Technology | #13969 | 1:1000 |
|  | Anti-H3K27me3 | Cell Signaling Technology | #9733 | 1:1000 |
|  | Anti-H3K36me3 | Cell Signaling Technology | #4909 | 1:1000 |
|  | Anti-H3K79me3 | Cell Signaling Technology | #74073 | 1:1000 |
|  | Anti-pan methyl lysine | Abcam | ab7315 | 1:2000 |
|  | Anti-total histone H3 | Proteintech | 17168-1-AP | 1:5000 |
|  | Anti-SETDB1 | Proteintech | 11231-1-AP | 1:2000 |
|  | Anti-SUV39h1 | Proteintech | 10574-1-AP | 1:2000 |
|  | Anti-EHMT2 | Proteintech | 66689-1-Ig | 1:5000 |
|  | Anti-GAPDH | Proteintech | 60004-1-Ig | 1:50000 |
| Secondary | HRP-conjugated anti-mouse IgG | Proteintech | SA00001-1 | 1:20000 |
|  | HRP-conjugated anti-rabbit IgG | Proteintech | SA00001-2 | 1:20000 |

